# Distribution of Sigma factors delineates segregation of virulent and avirulent *Mycobacterium*

**DOI:** 10.1101/235986

**Authors:** Aayatti Mallick Gupta, Sukhendu Mandal

## Abstract

Sigma factors, in combination with RNA polymerase and several transcription factors play specific role in expression of housekeeping as well as various stress responsive genes in mycobacterial species. The genus *Mycobacterium* includes a wide range of species under major pathogens, opportunists and non-pathogens. The number and combination of sigma factors is extremely diversified among *Mycobacterium*. We have performed comparative genome analysis among 40 different species of *Mycobacterium* whose whole genome sequence is available, in order to identify the distribution of sigma factors. The study illustrate that SigC, SigD, SigG, SigH, SigK and SigI are dominant among the true pathogens. Moreover, 16S rDNA based phylogenetic analyses distinctly differentiate the slow growing *Mycobacterium* from the fast growers, and clusters the true pathogens from the opportunists and non-pathogens. While evaluating the similarity coefficient upon the allotment of sigma factors of different *Mycobacterium* species through UPGMA dendrogram analysis, it is apparent that the true pathogens are grouped separately following the similar trend observed from evolutionary approach. Sigma factors playing dominant role in pathogenicity are found stable in nature with high aliphatic index thereby remain flexible at a wide range of temperature. The comparative distribution of six well known virulence factors of *Mycobacterium* - PhoP, PcaA, FbpA, Mce1B, KatG and PE_PGRS and various sigma factors justify the allotment pattern of mycobacterial sigma factors among pathogenic species. The pathogenicity responsible sigma factors elicit close resemblance with few notable characters of the known virulence factors. Thus the analysis renders that the distribution of sigma factors of different species of *Mycobacterium* can be a potential tool to predict the pathogenicity index of this genus.

## INTRODUCTION

Sigma factors are the extensive coordinator of transcription and have a typical role in transcription initiation through the recognition of specific promoter sequences of various set of genes. It is believed that changes in environmental factors lead to the replacement of sigma factors in the holoenzyme and the transcriptional regulation of a different set of genes. This in turn help in the expression of proteins that can help the organism to survive under stress conditions^1^. It is generally observed that every sigma factor recognizes distinct sets of promoter sequence. Therefore, variation in active sigma factor populations may represent a powerful way to modulate transcription profiles of an organism in accordance with its physiological requirements.

For having a complex physiological life style the species of *Mycobacterium*, as other actinomycetes, have been evolved in the past while present in the soil. However, this genus comprise of variety of strains that are associated with infectious diseases in a wide range of hosts. The major advancement of *Mycobacterium* species is an association of deletion (nonfunctional genes are deleted/inactivated and subsequently eroded) and insertion of genes (horizontal transfer and gene duplication) which enable their survival in various stresses^2-5^. The emergence of pathogens and opportunists from non-pathogens and vise-versa is an interesting study of investigation. Specially in presence of advancement of genomics which enrich nucleic acid databanks with whole genome sequence data. Pathogens often harbour chromosomal gene clusters encoding virulence functions, known as pathogenecity islands, which have been acquired by horizontal gene transfer, and allow such pathogens to infect the host^6-7^. Horizontal gene transfer indicate the addition of genetic elements transferred from the donor organism directly into the genome of the recipient organism, where they form genomic islands—that is, stretch of DNA which contain mobile genetic elements. Genomic islands may contain large chunks of virulence determinants (adhesins, invasins, toxins, protein secretion systems, antibiotic resistance mechanisms, etc), and thus are described as pathogenicity islands. Pathogenicity islands comprise of approximately 10–200 kilobases of genomic DNA that are unique in pathogenic bacterial strains but absent from the genomes of non-pathogens of the same or related species. Pathogenicity islands are supposed to have been procured as a block by horizontal gene transfer owing to (a) their G+C content is notably different from that of the genomes of the host micro-organism; (b) they are often flanked by direct repeats; (c) they are often integrated with tRNA genes; (d) they are associated with integrase determinants and other mobility loci; and (e) they demonstrate genetic instability. Indeed, all three mechanisms for genetic exchange or transfer between bacteria (that is, transformation, transduction, and conjugation) plays vital role in the evolution of pathogenic species^8^.

Hence, the expression of such gene clusters tent are acquired from various sources should depend on specialized transcriptional machinery which essentially includes different flavour of sigma factors with a constant set of core RNA polymerase. Thus it is an intriguing issue to find out the correlation between distribution of sigma factor and pathogenecity among mycobacterial species. Availability of whole genome sequences has opened the possibility to evaluate the degree of variation of sigma factor among the different species of *Mycobacterium.* Based on their pathogenecity index, the genus *Mycobacterium* can be grouped in pathogens, opportunists and non-pathogens ; involving both slow and rapid growers. In this study we include 40 Mycobacterial species in order to map their pathogenecity index as well as distribution of sigma factor.Among the 11 slow growing pathogensof this study, *M. tuberculosis*, *M. bovis*, *M. africanum*, *M canettii* and *M. microti* belongs to *M. tuberculosis* complex (MTBC)^9^, while the rest are included within non-tuberculous *Mycobacterium* (NTM). The study also comprises of 20 opportunists species of *Mycobacterium*, among them 12 belongs to slow growing variety and the rest 8 are rapid growers. Thus, opportunists show heterogeneous growth rate. They primarily belong to NTM group and are the causal agent of pulmonary and other disseminated infections in immunocompromised individuals^10^. The slow growing opportunists that belong to *M avium* complex (MAC) is formed by *M avium*, *M intracellularae* and *M colombiense*^11^. A recently described *M yongonense* also belongs to MAC, and phylogenetically related to *M intracellularae*^12^. *M vulneris* recently individualised among MAC, previously referred to as *M. avium* sequevar Q is closely related to *M. colombiense*^13^. M *triplex* closely resembles MAC in biochemical tests but failed to react with the commercial probe designed for MAC^14^. *M. tusciae* is a slow growing opportunist isolated from the lymph node of an immunocompromised child. It shows evolutionary proximity to the fast-growing forms^15^. *M. haemophilum* is a slow growing ‘blood-loving’ *Mycobacterium* prefers to grow at low temperature range. It frequently causes skin infection to immunocompromised patients. Close genetic relatedness of *M. haemophilum* is found with *M. ulcerans* and *M. marinum* whereas in regard to fatty acid composition there exist an interesting similarity between *M. haemophilum* and *M. Leprae*^16^. The study includes *M. indicus pranii* as the only slow growing non-pathogen that belongs to MAC^17^.

The opportunists that includes rapid growing *Mycobacterium* (RGM) are *M. abscessus*, *M. neoaurum, M. fortuitum*, *M. thermoresistibile*, *M. cosmeticum*, *M. mageritense*, *Mgoodii* and *M chelonae*, causing infections in immunodeficient patients. The study delineates with 8 different non pathogenic species of RGM. *M vaccae* is a soil *Mycobacterium*, functions as an antidepressant as it stimulates the generation of serotonin and nor-epinephrine in the brain^18^. *M. vanbaalenii* is a free-living RGM that utilises polycyclic aromatic hydrocarbons (PAH), closely related to *M. vaccae*^19^. *M. hassiacum* is RGM and thermophilic in nature. It shows a high level of similarity with the slow growing *M. xenop*i^20^. *M. phlei, M. rhodesiae*, *M. chubuense* and M. *smegmatis* are the other non-pathogenic species of *Mycobacterium*^21-22^ included in this study.

The present work identifies distribution of the sigma factors among slow and rapid growing *Mycobacterium*, of 40 different species of *Mycobacterium* that includes pathogens, opportunists and non-pathogens. Remarkably, the distribution of sigma factors ascertains its role in pathogenicity. These analyses provide strong evidence of a key role played by sigma factors in pathogenesis of different species of *Mycobacterium.*

## RESULTS

### 16S rRNA phylogeny segregrates pathogenic and non-pathogenic Mycobacterium

To infer the evolutionary relationships of pathogens, opportunists and non-pathogens among the 40 different species of *Mycobacterium*, a comprehensive phylogenetic tree is constructed considering 16S nucleotide sequence. The neighbour-joining (NJ) tree built for this dataset based upon the linear sequence of 16S rDNA is shown in supplementary figure S1 and that of 16S secondary structure based phylogeny is elicited in figure 1. Overall, both the analysis concatenates to form a distinct clade between the slow growing and the rapid growing species of *Mycobacterium.* Furthermore, the trees have well fulfilled to demarcate pathogens, opportunists and non-pathogens. Slow growing pathogenic forms are distinguishable from slow growing opportunists with an exception of *M. farcinogens* which itself is a slow grower but shows evolutionary resemblance with rapid growers^23^. Similarly, the slow grower *M. tusciae* shows affinity with RGM^15^. Evidence from the evolutionary approach establishes the fact that rapid growing forms of opportunists and non-pathogenic *Mycobacterium* are not distinctly distinguishable from each other. *M. hassiacum* offers a peculiarity for being ancestrally close to the slow-growing *M. xenopi*^17^. The study reflects the underlying fact of the relationship between growth rate and pathogenicity among different *Mycobacterium.* All pathogenic varieties are ubiquitously slow growers. The opportunists are diversely populated belonging to both the slow and the rapid growing variety of *Mycobacterium.* The non-pathogens are basically rapid growing form with an exception of *M. indicus pranii.* It is a non-pathogen and belongs to MAC. The rapid growing pathogenic variety of *Mycobacterium* is not found anywhere from the present analysis. Hence, the study provokes the fact that as the growth rate increases pathogenicity diminishes showing an inverse relationship between the two. Pathogens comprise of the slow growing while, the non-pathogens consist of the rapid growing *Mycobacterium.* The opportunists are intermediate in position and are most diversified forms.

**Figure 1:**
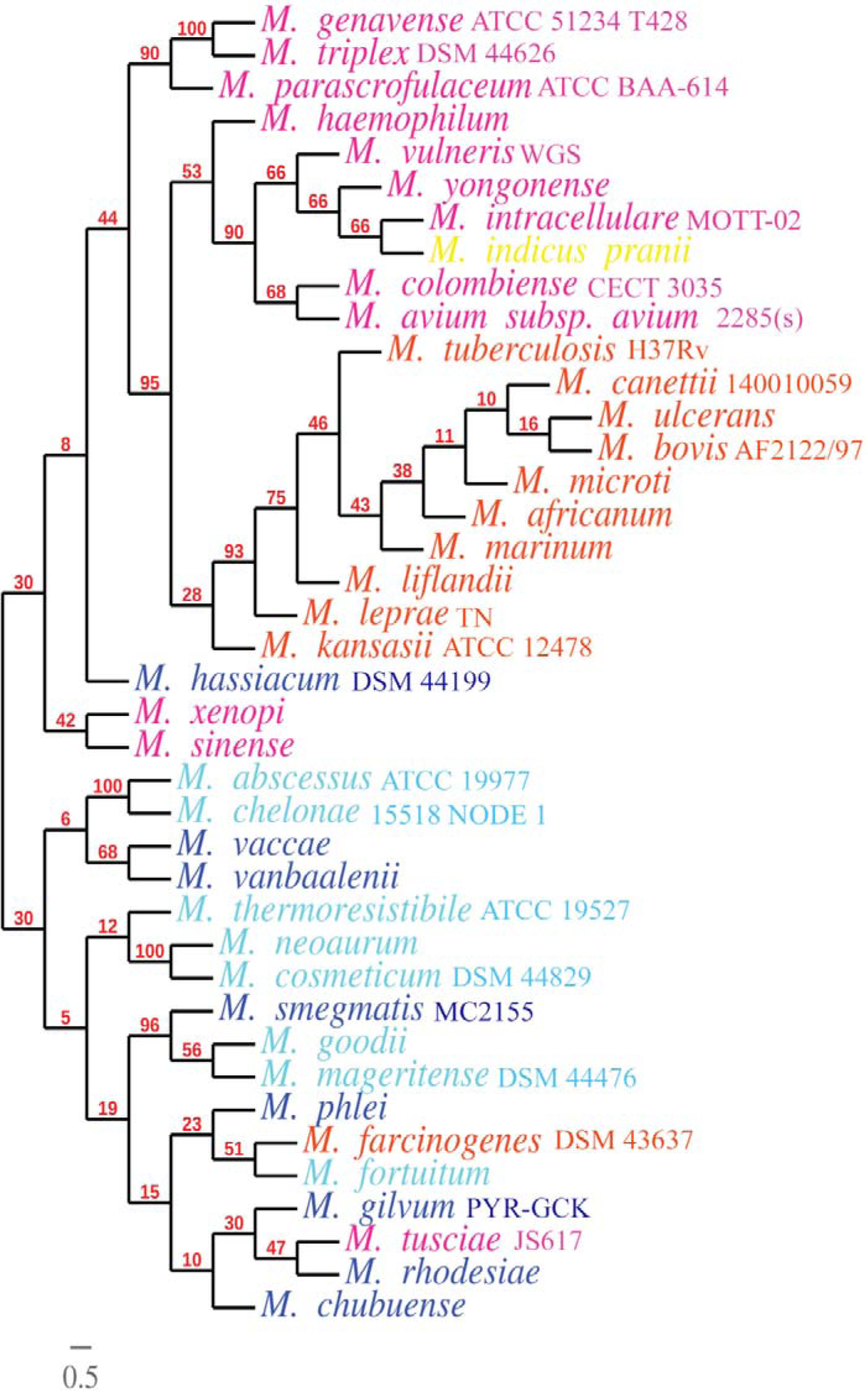
Phylogenetic tree showing the relationships among different species of *Mycobacterium* based on secondary structure of 16S rRNA. Slow growing pathogen is demarcated in orange, opportunists that are slow grower in crimson and the only slow growing non-pathogen in yellow. Rapid grower non-pathogen is marked in blue while opportunists that are rapid grower are indicated in cyan.

### The distribution of sigma factors in Mycobacterial species follows their phylogenic grouping

In this study, the distribution of sigma factors of *M. tuberculosis* is optimised with that of 40 different species of *Mycobacterium.* On evaluating the similarity matrix with Jaccard’s coefficient for UPGMA dendrogram analysis (figure 2) it is apparent that pathogenic forms are grouped separately following the evolutionary trend observed in phylogenetic analysis. Nevertheless, pathogens are grouped separately from that of opportunists and non-pathogens in the dendrogram analysis while the slow growers and rapid growers are found to merge together. This analysis of similarity coefficient upon the arrangement of sigma factors renders similar cluster pattern in terms of virulence (not on the growth rate) with respect to the evolutionary trend. Thus, sigma factor can be an essential tool to demonstrate the differential virulence pattern in various species of *Mycobacterium.*

**Figure 2:**
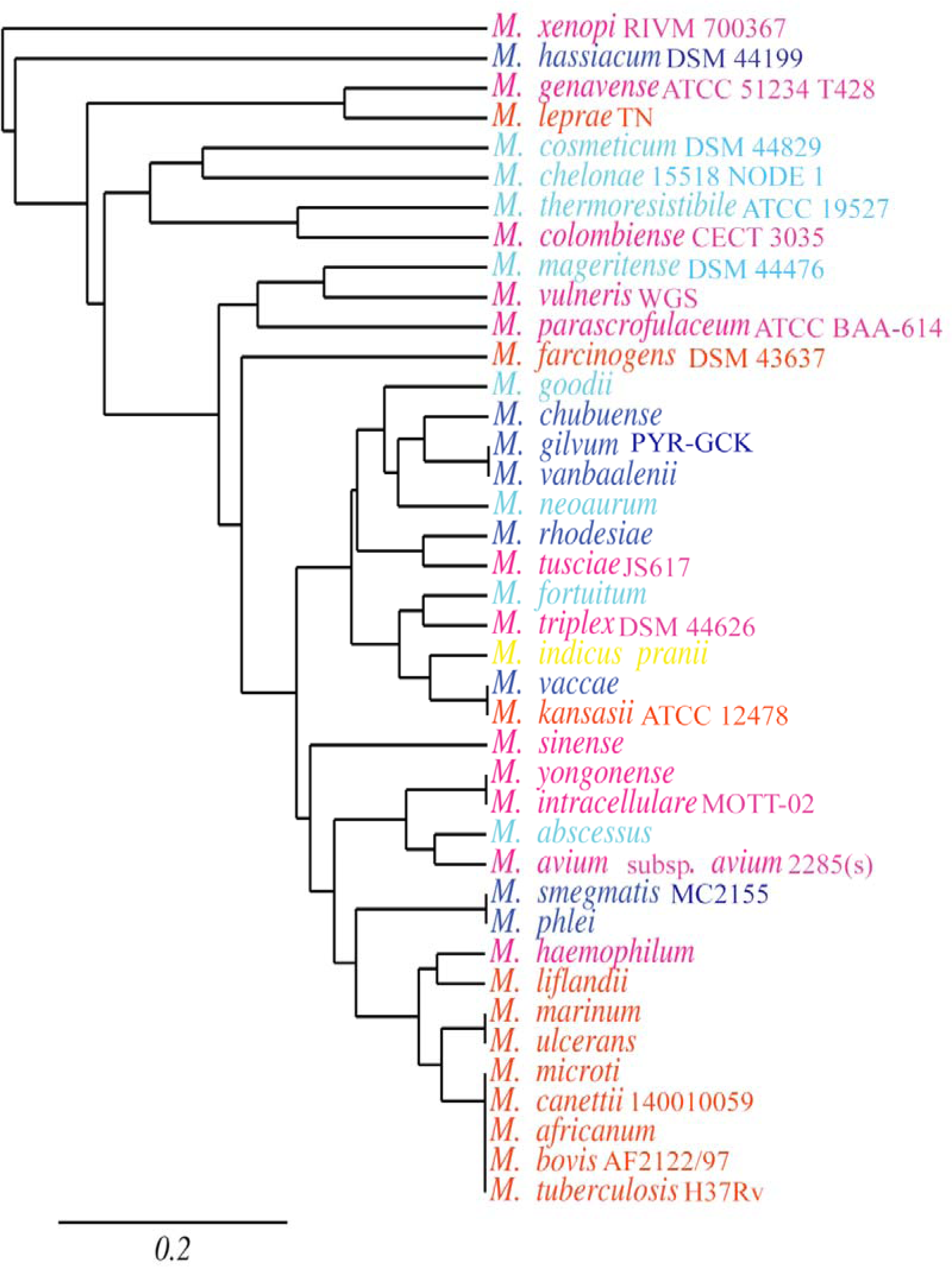
Dendrogram derived from cluster analysis (UPGMA) using the Jaccard’s similarity coefficient based on the distribution of various sigma factors of different species of *Mycobacterium.* Colour demarcation is same as that in figure 1.

### Occurence of sigma factor in *Mycobacterium* is a potential tool to predict their pathogenicity index

Based upon the percentage of occurrence of the sigma factors among 40 different *Mycobacterium* species it is evident that the ECF sigma factors - SigC, SigD, SigG, SigH, SigK and SigI are widely found among pathogens than that in opportunists and nonpathogens (Table 1). SigC shows 90.9% occurrence among pathogens, 20% among opportunists while only 11.11% among non-pathogens. SigD is existed at 81.81% among pathogens, while only 40% and 33.33% is found among opportunists and non-pathogens respectively. This scenario is similar in case of SigG, SigH, SigK and SigI indicating their dominant role on pathogens (Figure 3). However, primary sigma factor SigA, primary-like sigma factor SigB and the alternative sigma factor SigF are found to be equally distributed in all the 40 different species of *Mycobacterium* chosen for the present study. Thus it depicts their indispensable function as housekeeping genes in contrast to ECF sigma factors that are adapted to specific environmental conditions. The study ascertains that the other ECF sigma factors like SigE, SigJ and SigM have shown their omnipresence allocation among 40 different species of *Mycobacterium.*

**Figure 3:**
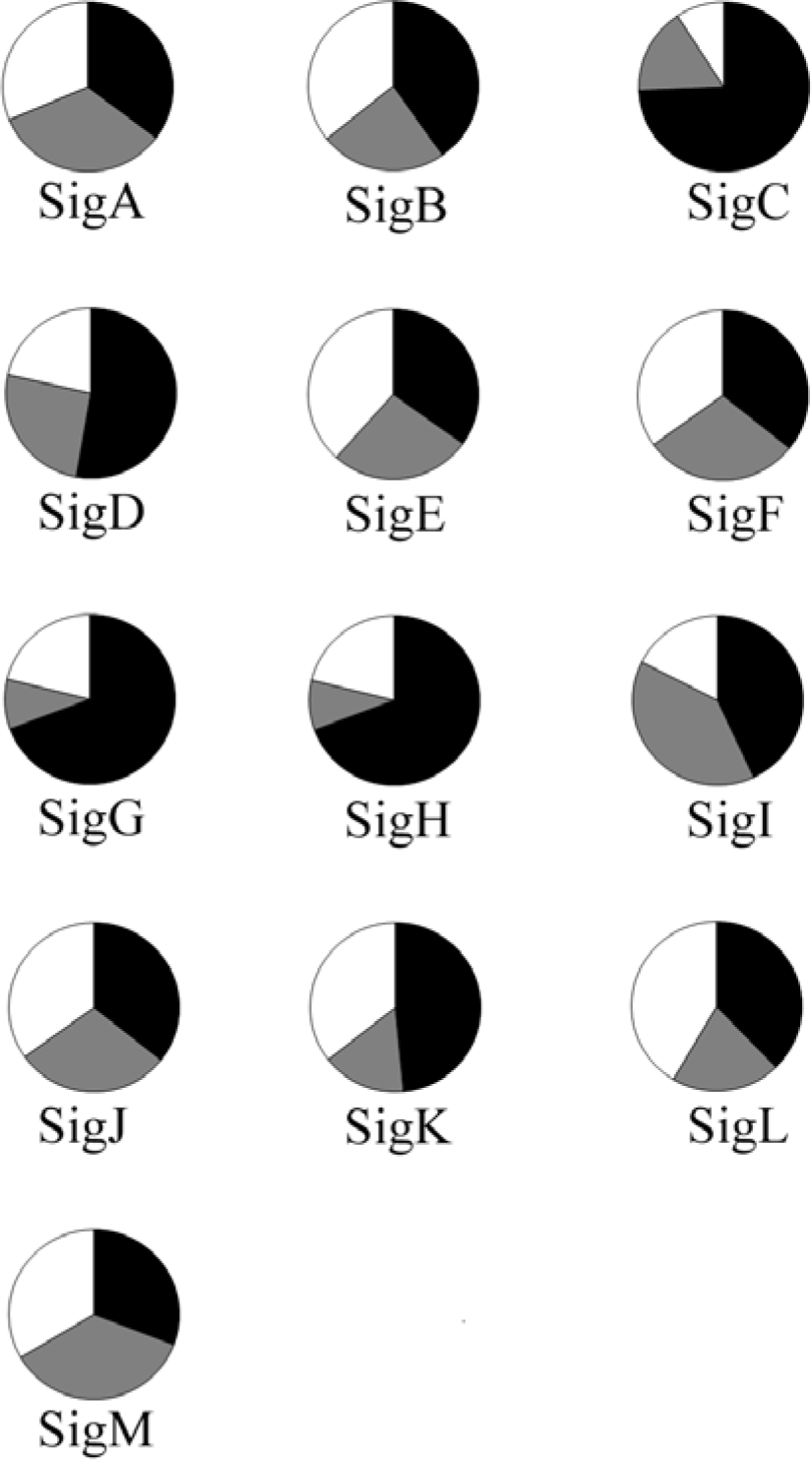
Pie-chart showing the percentage of occurrence of each sigma factor in different mycobacterial species. The colour demarcations are as pathogens-black, opportunists-grey and non-pathogens-white. Figure 4: *In silico* physio chemical analysis of the sigma factors. (a) Instability index of sigma factors of *M. canettii 140010059* (representative of pathogens) in black, *M. yongonense* (representative of opportunists) in grey and *M. phlei* (representative of non-pathogens) in white. Instability index > 40 is indicative of unstable protein while < 40 means the protein is stable (demarcated with black line). (b) Aliphatic index (AI) of sigma factors of *M. canettii 140010059* (representative of pathogen) in black, *M. yongonense* (representative of opportunist) in grey and *M. phlei* (representative of non-pathogen) in white. . (c) GRAVY value indicates sigma factors in the various species of *Mycobacterium* ..

**Fig.4:**
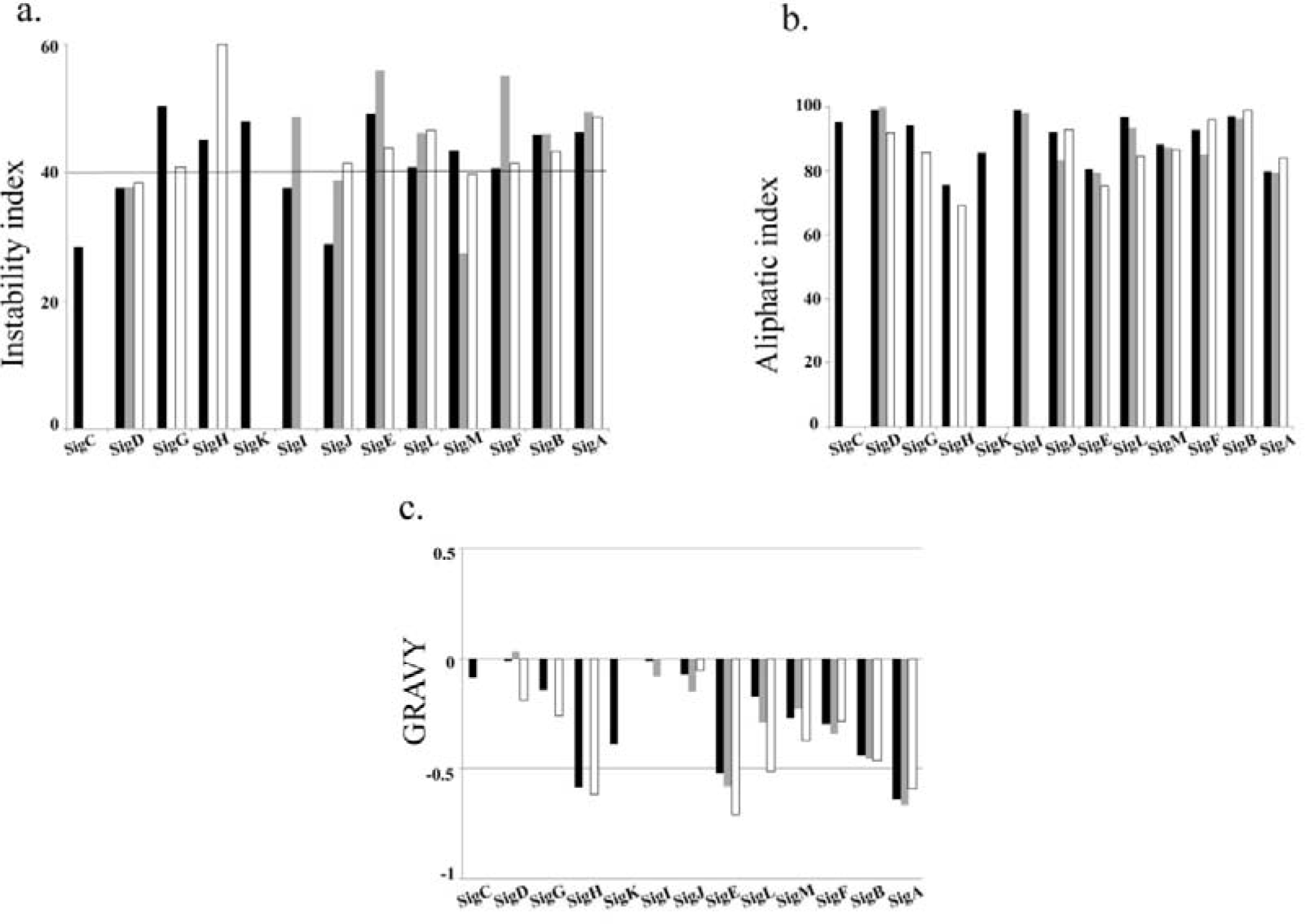

**Table1:** Distribution of various sigma factors among pathogens, opportunists and non-pathogens in different species of *Mycobacterium*.

| Sigma factor | Pathogens | Opportunists | Non-pathogens |
| --- | --- | --- | --- |
| SigC | 90.90% | 20.00% | 11.11% |
| SigG | 72.72% | 40.00% | 22.22% |
| SigH | 72.72% | 5.00% | 22.22% |
| SigI | 54.54% | 50.00% | 22.22% |
| SigD | 81.81% | 40.00% | 33.33% |
| SigK | 90.90% | 30.00% | 66.66% |
| SigJ | 90.90% | 75.00% | 88.88% |
| SigM | 81.81% | 95.00% | 88.88% |
| SigF | 90.90% | 75.00% | 88.88% |
| SigB | 100.00% | 60.00% | 88.88% |
| SigA | 100.00% | 95.00% | 88.88% |
| SigE | 90.90% | 70.00% | 100.00% |
| SigL | 90.90% | 50.00% | 100.00% |

### Comparative pathogenomics study of *Mycobacterium*

The 6 well known virulence factors of *Mycobacterium* is chosen for the present study (generated against VFDB: refer the methods section), which includes (a) proteins for cell wall biosynthesis – FbpA, PcaA; (b) mammalian cell entry protein – Mce1B; (c) protein related to stress adaptation – KatG; (d) a regulatory protein that senses Mg^2+^ starvation and controls expression of virulence responsive genes – PhoP and (e) a PE family protein exclusively found in the genus *Mycobacterium* – PE_PGRS. FbpA is a fibronectin binding protein that enhances the uptake of *Mycobacterium* onto macrophages via complement-mediated phagocytosis. It is related to mycolyltransferase activity that transfers long-chain mycolic acids to trehalose derivatives pivotal for the biosynthesis of the Mycobacterial cell wall and for the survival of *Mycobacterium*^24-25^. The cell wall protein, PcaA acts as cyclopropane synthase that incorporates a single proximal cycloproprane ring on the α-mycolic acids and the production of cord factor in the cell wall required for persistence and virulence^26-27^. Mce1B belongs to a mce family protein playing an essential role in bacterial virulence imposing their role at the route of infection^28^. KatG is a stress-responsive protein that plays major role in the degradation of catalase:peroxidise generated by phagocyte NADPH oxidase^29^. The regulatory protein PhoP senses Mg^2+^ starvation which controls the expression of genes involved in surface remodelling and adaptation to intracellular growth^30^. PE_PGRS belongs to the PE family of protein consisting of polymorphic GC-rich repetitive sequence at its C-terminal domain that inhibits proteasomal degradation of the N-terminal PE domain^31^ thus inhibiting antigen processing by CD8+ T cells.

While analysing the distribution pattern of these well known virulence factors among the 40 different species of *Mycobacterium*, it is found from the UPGMA dendrogram based similarity coefficient that these virulence factors follows similar cluster pattern that is observed in case of allotment of the sigma factors (figure S2). The Pathogens are well differentiated from opportunists and non-pathogens. All the virulence responsive proteins are widely populated among pathogens than that is present among the opportunists and non-pathogens (figure S2). FbpA exhibits their 100% occurrence among pathogens, 35% in opportunists and 44.44% in non-pathogens (Table 2). It is interesting to note that PE_PGRS is exclusively found among the pathogens. Thus the trend observed in sigma factors is followed for other virulence factors as well.

**Table 2:** Distribution of 6 well known virulence factors among pathogens, opportunists and nonpathogens in *Mycobacterium*

| Virulence factors | Pathogens | Opportunists | Non-pathogens |
| --- | --- | --- | --- |
| Mce1B | 81.81% | 15.00% | 22.22% |
| PcaA | 90.90% | 15.00% | 33.33% |
| FbpA | 100.00% | 35.00% | 44.44% |
| PhoP | 90.90% | 35.00% | 66.66% |
| KatG | 100% | 55% | 77.77% |
| PE_PGRS | 90.90% | - | - |

### Pathogenecity responsive sigma factors follow the character of pathogenecity responsive factors

The aliphatic index (AI) of a protein describes the comparative volume utilised by aliphatic side chains (alanine, valine, isoleucine, and leucine). It is considered as a positive factor for enhancing the thermostability of globular proteins^32^. Overall the AI for the sigma factors ranged from 70.69-108.62. Particularly, SigC, SigD, SigG, SigK, and SigI have shown a very high AI than the rest indicating that these sigma factors may be stable for a wide range of temperature. The result thus interprets the fact that the sigma factors which show their wide existence among pathogens have high AI depicting an increase in thermal stability.

The Grand Average hydropathy (GRAVY) value for a peptide or protein is calculated as the sum of hydropathy values of all the amino acids, divided by the number of residues in the sequence^33^. GRAVY demonstrates solubility of a protein where hydrophobicity corresponds to a positive value and hydrophilicity corresponds to a negative value. GRAVY indices for sigma factors of 40 different species of *Mycobacterium* exhibit hydrophilicity in majority of the cases while a few of them shows hydrophobicity. The lower range of value elicits its greater extent of hydrogen bonding with water molecules and thus higher is its solubility. These analyses reveal that SigD of *M. avium* subsp. *avium* 2285(s), *M. intracellularae* M0TT-02, *M. yongonense* and *M. indicus pranii* are hydrophobic in nature. SigI of *M. tuberculosis* H37Rv, *M. bovis* AF2122/97, *M. microti, M. farcinogens* DSM 43637, *M. tusciae* JS617 and *M. abscessus* ATCC 19977 renders hydrophobicity. In SigJ the organisms that show hydrophobicity includes *M. ulcerans*, *M. liflandii*, *M. marinum*, *M. haemophilum*, *M. vanbaalenii*, *M. gilvum* PYR-GCK and *M. chubuense.* The instability index (II) evaluates the stability of the protein under *in vitro* conditions. A protein whose II is below 40 are predicted as stable while a value above 40 elicits that the protein may be unstable. The instability of proteins is possibly determined by the order of certain amino acids in its sequence in accordance with the presence of certain dipeptides occurring differently in unstable and stable proteins^34^. SigC, SigD, SigI and SigJ of majority of *Mycobacterium* are stable in nature while instability is observed for most of the rest sigma factors. An example of AI, GRAVY and II analysis of sigma factors with each representative from pathogens ( *M. canettii140010059*), opportunists ( *M. yongonense*) and non-pathogens ( *M. phlei*) is illustrated in figure 4.

In case of 6 well known virulence factors chosen for the study, AI ranges from 63.58-106.78. The mammalian cell entry protein Mce1B and PE family protein PE_PGRS estimates a very high AI ensuring stability over a wide range of temperature (Supplementary figure S4-a). GRAVY value shows the dominance of hydrophilicity among the virulence factors except for PE_PGRS, which manifests predominance in hydrophobicity (Supplementary figure S4-b). II of PhoP is mostly unstable in nature excepting the ones belonging to Mtb complex. Overall the virulence factors indicate that most of the *Mycobacterium* is stable in nature (Supplementary figure S4-c).

## DISCUSSION

In this study, the distributions of sigma factors of 40 different species of *Mycobacterium* are compared. The chosen set encompasses pathogens, opportunists and non-pathogens from both slow and rapid growing species of *Mycobacterium.* The varied growth rate and pathogenicity of these organisms along with the number of sigma factors they bear are interlinked in table 3. From the analysis it has been found that all pathogens are included only within the slow growing variety while non-pathogens are typically rapid growers. Opportunists belong to both the slow and rapidly growing variety. Thus, a close correlation is derived between the growth rate and pathogenicity portraying that the intensity of virulence diminishes from slow growing to rapid growing *Mycobacterium.* Based on 16S rDNA sequence, the phylogenetic analysis of the 40 various species of *Mycobacterium*, efficiently discriminates the slow growers from the rapid growers. Likewisethe evolutionary analysis based upon the primary sequence of 16S rDNA as well as those from the secondary structure annotated phylogenetic tree prominently discriminates pathogens from opportunists and nonpathogens. These finding flash enough light on the distribution of pathogenecity trend within *Mycobacterium* species. The gaining of the pathogenecity trait happened in cost of their growth rate. It is an intriguing question why the pathogens need to be a slow grower *in vitro* and *in vivo.* It might be true that pathogens are dependent on some host factor for their proliferation; however it is unlikely to be the only cause as the growth rate does not change while supplemented with host factors. In this point it is demanding to explore the truth that lies in gain of pathogenecity in cost of growth rate or loss of pathogenecity to increase it. Similar study^35^ based on comparative genome analysis of *Mycobacterium* was successful to understand the genome feature of each pathogenic and non-pathogenic species based on 16S rDNA to its unique niche. However the position of opportunists has not been considered earlier. It has been noted from the present study that slow growing true pathogens are distinctly distinguishable from slow growing opportunists, while, the opportunists that are rapidly growing form are found in combination with rapid growing non-pathogens. Thus, opportunists are assembled variedly onto the slow and rapid growers. Moreover, the evolutionary lineage of opportunists justifies its diversification from gaining pathogenecity.

**Table 3:**
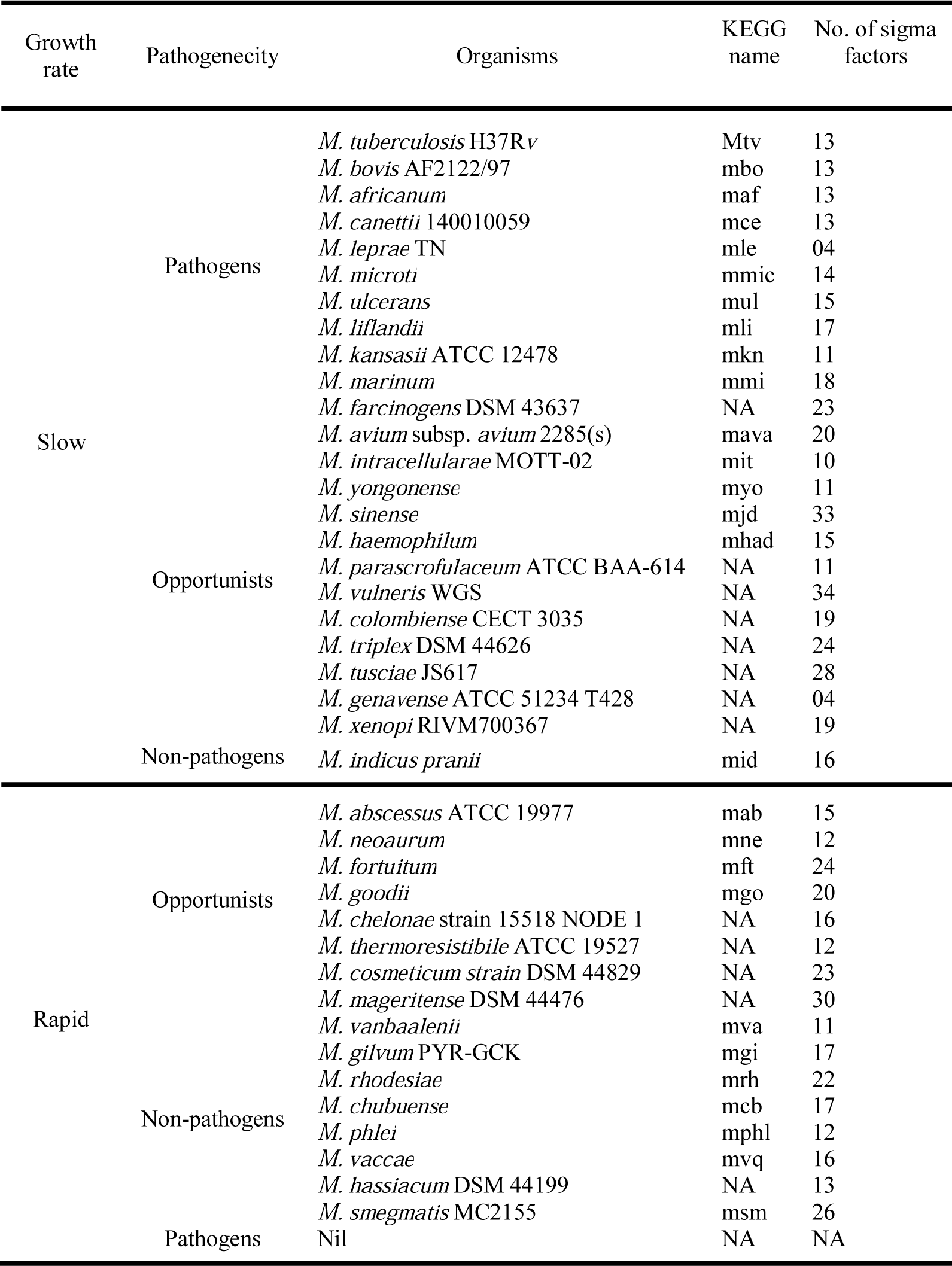
List of studied Mycobacterial species with their growth rate, pathogenecity and number of sigma factors present in these.

The study is emphasised on the overall difference among pathogens, opportunists and non-pathogens based on the distribution of sigma factors such as to determine the role of a particular sigma factor on pathogenicity. Jaccard’s similarity coefficient analysis based on sigma factor availability follows the evolutionary trend in terms of virulence. However, the distinction in growth rate pattern is absent in this UPGMA analysis. The discrimination of slow growers from the rapid growers observed during phylogenetic analysis is lost, yet the differentiation of pathogens from opportunists and non-pathogens is well maintained. The result is further affirmed with the analysis based on the investigation of the known virulence factors – PhoP, FbpA, PcaA, Mce1B and PE_PGRS on the 40 different species of *Mycobacterium.* The ECF sigma factors - SigC, SigD SigG, SigH and SigK is chiefly found to exist among pathogens signifying their key role to measure pathogenicity index. Besides, the principle sigma factor – SigA, primary like sigma factor – SigB and alternative sigma factor - SigF are uniformly distributed in all the *Mycobacterium* species included in this study. Among the ECF sigma factors, it is apparent that SigE, SigJ and SigM have exhibited universal existence in different species of *Mycobacterium*. The distribution of 6 other well known virulence factors included for this study manifested similar trend that is found in case of sigma factors prevalent among pathogens. Thus, from the present analysis; it might be helpful to predict the role of particular sigma factor in pathogenicity.

On computing the various physio-chemical properties of the sigma factors it is established that SigC, SigD, SigG, SigK, SigH, SigI of pathogens shows a very high AI, accountable for elevation in thermal stability. This is equally true in case of AI of the known virulent factors chosen for the study. A recent study on comparative proteomic analysis on two different strains of *M. tuberculosis* H37Rv (virulent) and H37Ra (avirulent) on PE/PPE multigene family reveals a transition from hydrophilicity to hydrophobicity. It has shown that certain PE family of protein is hydrophobic in H37Rv whereas its counterpart is hydrophilic in H_37_Ra^36^. This implicates that the virulent strain of M *tuberculosis* H37Rv is influenced towards hydrophobicity. Our study imparts that, despite the fact, that most of the sigma factors in the various species of *Mycobacterium* are hydrophilic in nature, certain sigma factors like, SigD and SigJ of NTM along with few SigI belonging to MTB complex contribute in hydrophobicity. While investigating II, it is elicited that the sigma factors which are widely prevalent among pathogens are relatively stable in nature. However, rest of the sigma factors broadly persisted in opportunists and non-pathogens are somewhat unstable in nature. This interpretation is moreover verified with II of the known virulence factors.

In summary, identifying sigma factors among 40 different species of *Mycobacterium* comprising the slow and rapid grower as well as pathogens, opportunists and non-pathogens, it is evident that sigma factors can be a potential tool to predict pathogenicity. The analysis focuses upon the propagation of different sigma factors upon the diverged category of *Mycobacterium* species.

## Methods

### Retrieval of nucleotide and amino acid sequence of Mycobacterial sigma factors

The sigma factors of different *Mycobacterium* species have been obtained using the advance search mode of uniprot^37^ and KEGG orthology database search (http://www.kegg.jp/ or http://www.genome.jp/kegg/)^38-40^.

### Phylogenetic analysis based on primary sequence of 16S rDNA

The 16S rDNA sequences of 40 different species of *Mycobacterium* used in the study have been retrieved from NCBI database. MEGA 6^41^ is used for sequence-based tree construction with progressive multiple sequence alignment (MSA) algorithms i.e., CLUSTRALW (inbuilt in MEGA 6) followed by the test of phylogeny inferred with neighbour-joining (NJ) method along with kimura 2 parameter model as distance correlation. In order to test the reliability of the tree branches, a bootstrap analysis with 1000 replicates is performed.

### Phylogenetic analysis based on secondary structure of 16S rDNA

MAFT version 7^42^ is employed for the test of phylogeny taking into account the secondary structure of 16S rDNA of 40 different species of *Mycobacterium* used for the study. Q-INS-i of MAFT programme utilises the Four-way consistency objective function for incorporating structural information. The structure annotated phylogenetic tree file in ‘NEWICK’ format is generated by MAFT version 7 which offers building of phylogenetic tree using TreeDyn 189.3^43^ The inclusion of secondary structure information provides a robust analysis that incorporates additional biological information that strengthens the confidence that positional homology is being conserved.

### Statistical data analysis

Binary data based on the distribution of different sigma factors as well as other virulence factors in Mtb and rest other species of *Mycobacterium* have been analysed by Jaccard similarity coefficient^44^. The similarity matrix thus obtained is further subjected for cluster analysis by the Unweighted Pair Group Method with Arithmetic Mean (UPGMA) method^45^. It is a simple agglomerative hierarchical clustering method to produce a dendrogram from a distance matrix. This method employs a sequential clustering algorithm, in which local topological relationships are inferred in order of decreasing similarity and a dendrogram is built in a stepwise manner. This study is helpful to generate the relatedness among the different species of *Mycobacterium*. The robustness of the nodes of dendrogram is tested by bootstrap analysis using 1000 resamplings. These entire analyses have been carried out using Dendro UPGMA web server (http://genomes.urv.es/UPGMA/). UPGMA dendrogram is drawn using TreeDyn 189.3^43^

### *Insilico* proteomics study

These include the comparison of instability index (II), aliphatic index (AI) and grand average of hydrophobicity (GRAVY) of the sigma factors and other well-known virulence factors in different species of *Mycobacterium.* It is carried out with the help of ProtParam tool from ExPASy portal (http://web.expasy.org/protparam/). ProtParam computes various physio-chemical properties deduced from a protein sequence. No additional information is required about the protein under consideration. The parameters analysed by ProtParam include the molecular weight, theoretical pI, amino acid composition, atomic composition, extinction coefficient, estimated half life, instability index, aliphatic index and grand average of hydrophobicity.

### Comparative pathogenomics analysis

These is harnessed using the virulence factor database (VFDB)^46^ that have allowed for the spontaneous comparison of 6 virulence factors among the different species of *Mycobacterium* used for the study. VFDB is an integrated and comprehensive online recourse for curating information about virulence factors of bacterial pathogens. It provides a solid platform of the best characterised bacterial pathogens, with their structural features, functions and mechanisms adapted to conquer new niches and to arrest host defence mechanism, to cause disease. This database aims to develop innovative rational approaches in the eradication of the infectious diseases.

## Acknowledgement

Fellowship of AMG and part of the research is supported by grant no 548 (Sanc)/ST/P/S&T/9G-5/2015 funded by Department of Science &Technology, Govt. of West Bengal, India. The authors are thankful to the High Performance Computing for Modern Biology, University of Calcutta, to provide the infrastructure to carry forward the work.

## Conflict of interest statement

None declared.

## Author contributions

AMG did the experiment and analysed the data. SM designed the experiment, interpreted the data and prepared the manuscript.

---

**Figure S1:**
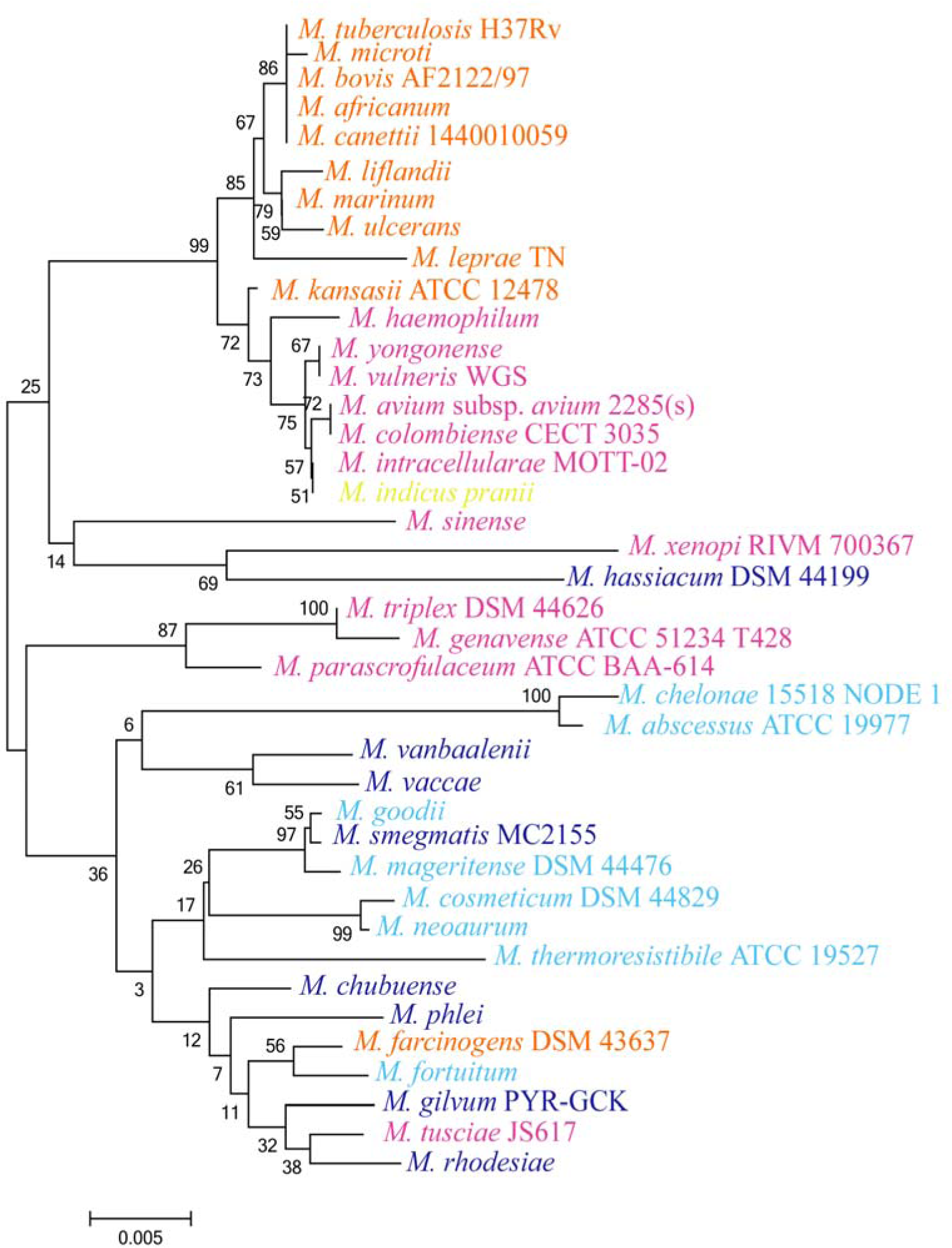
Phylogenetic tree showing the relationships among the 16S rDNA sequences of different species of *Mycobacterium* based on nucleotide sequence alignment using neighbour-joining method. Bootstrap values are calculated from 1000 replications of Kimura 2-parameter. (Bar = 0.002 nucleotide substitution per position). Colour demarcation is same as in figure 1.

**Figure S2:**
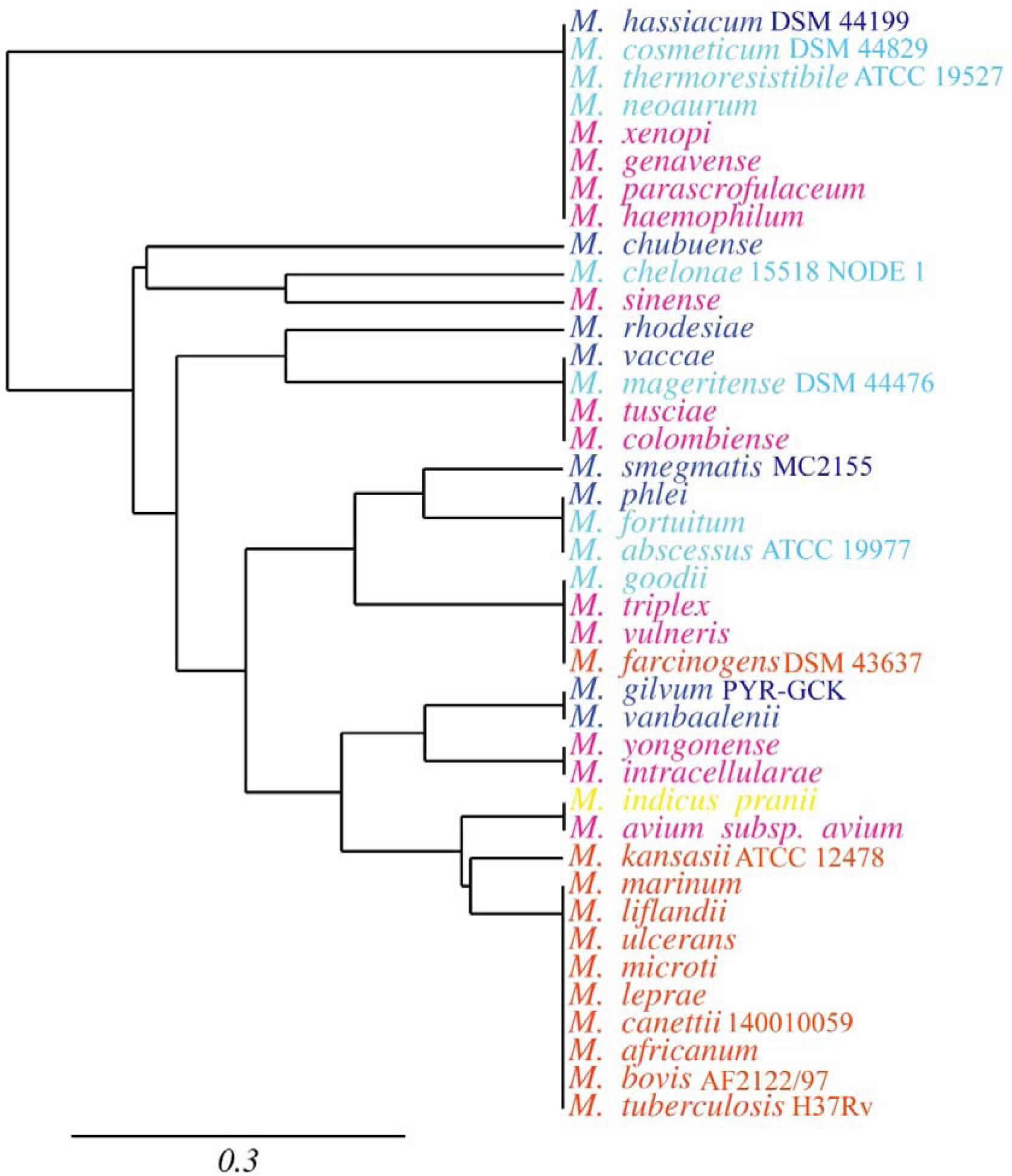
Dendrogram derived from cluster analysis (UPGMA) using the Jaccard’s similarity coefficient based on the distribution of virulence factors of different species of *Mycobacterium.* Colour demarcation is same as in figure 1.

**Figure S3:**
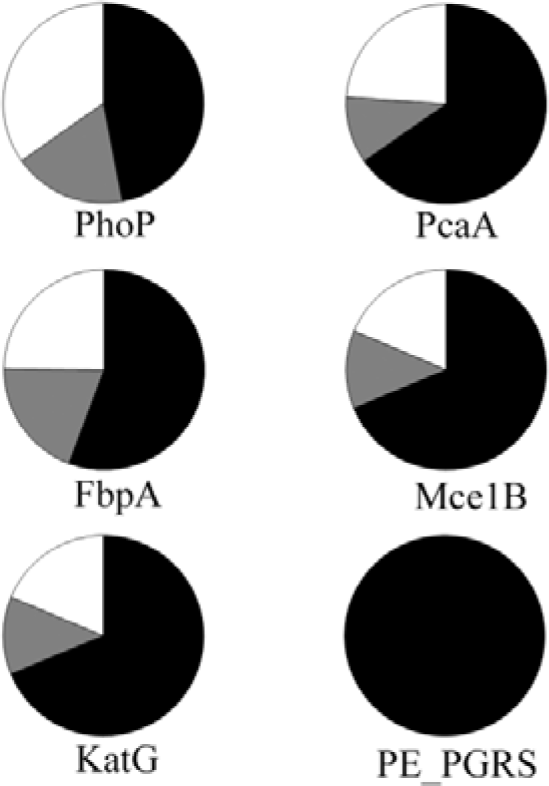
Pie-chart showing the percentage of occurrence of 6 well known virulence factors of *Mycobacterium.* The colour distinctions are followed from figure 3. Figure S4: *In silico* physio chemical study of 6 well known virulence factors of *Mycobacterium*. (a) Instability index of the different virulence factors of *M. canetti i140010059* (representative of pathogen), *M. yongonense* (representative of opportunist) and *M. phlei* (representative of non-pathogen). The colour demarcation is followed from figure 4. Instability index > 40 is indicative of unstable protein while < 40 means the protein is stable (demarcated with black line). . (b) Aliphatic index (AI) of all the virulence factors of *M. canettii 140010059, M. yongonense* and *M. phlei*, representing pathogen, opportunist and non-pathogen respectively. . (c) GRAVY value indicates virulence factors.

**Fig.S4:**
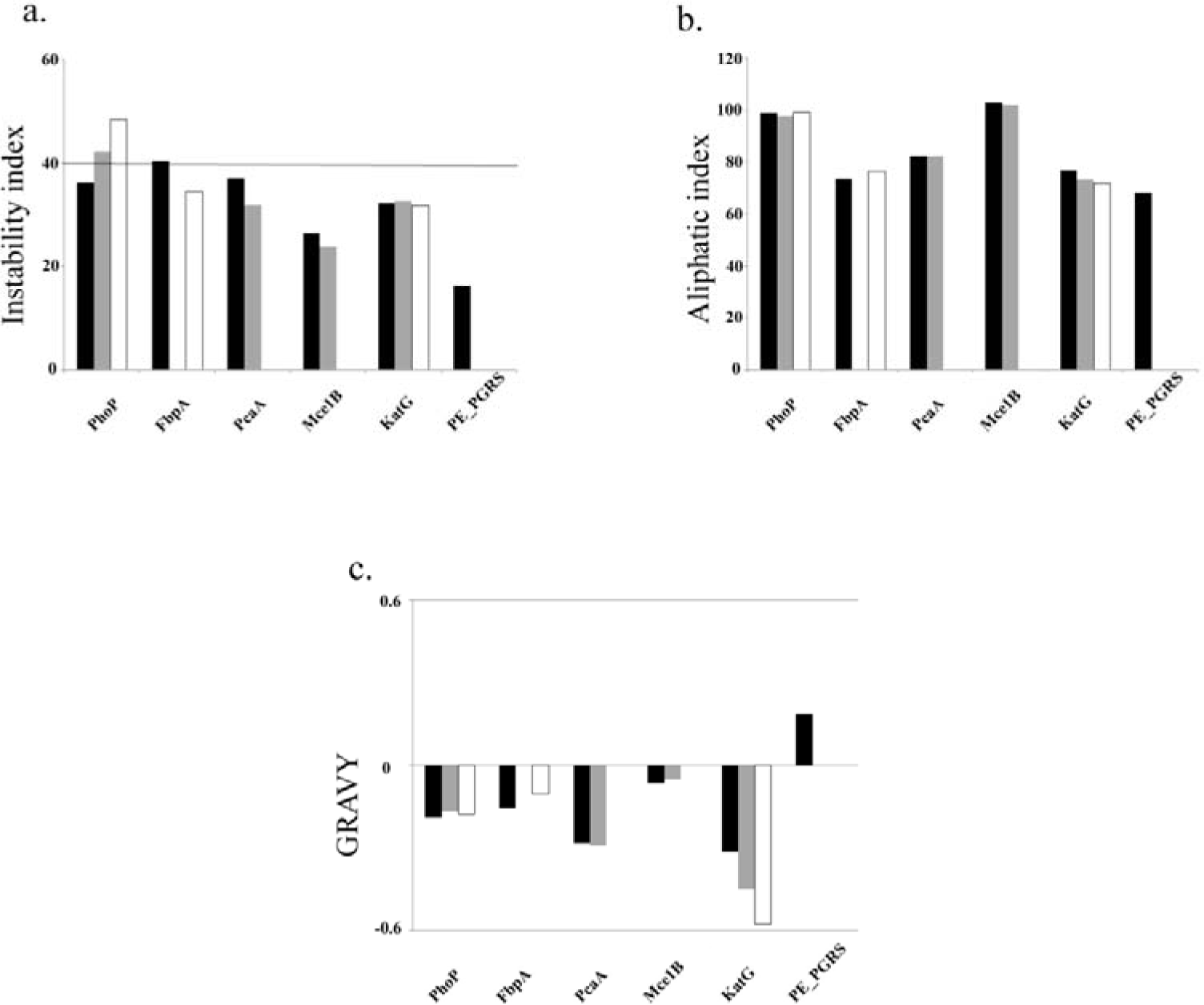

